# Motor Imagery Enhances Performance Beyond the Imagined Action

**DOI:** 10.1101/2024.10.04.616651

**Authors:** M. Gippert, P.-C. Shih, T. Heed, I. S. Howard, M. Jamshidi Idaji, A. Villringer, B. Sehm, V. V. Nikulin

## Abstract

Motor imagery is frequently utilized to improve the performance of specific target movements in sports and rehabilitation. In this study, we show that motor imagery can facilitate learning of not only the imagined target movements but also sequentially linked overt movements. Hybrid sequences comprising imagined and physically executed segments allowed participants to learn specific movement characteristics of the executed segments when they were consistently associated with specific imagined segments. Electrophysiological recordings revealed that the degree of event-related synchronization in the alpha and beta bands during a basic motor imagery task was correlated with imagery-evoked motor learning. Thus, both behavioral and neural evidence indicate that motor imagery’s benefits extend beyond the imagined movements, improving performance in linked overt movements. This provides new evidence for the functional equivalence of imagined and overt movements and suggests new applications for imagery in sports and rehabilitation.

## 1 Introduction

Motor imagery refers to the mental simulation, or rehearsal, of movement without physical execution. Executed and imagined movements exhibit strong similarities in timing ^1^, associated cardiac and respiratory activity ^2,3^, activated brain structures ^4^ and neural responses ^5^. This suggests shared mechanisms between overt and imagined movements ^6,7^. In addition, motor imagery supports learning during laboratory motor tasks such as sequence learning ^8^, strength ^9^ and balance tasks ^10^; in rehabilitation after neural ^11^ and musculoskeletal injury ^12^; and in sports performance improvement interventions ^13^. Even motor adaptation, the gradual adjustment of movements to changed environmental dynamics, benefits from motor imagery ^14^. Thus, motor imagery is an effective intervention for performance improvement across a wide variety of movement-related contexts (for meta-analysis see ^15^; for reviews see ^16^; ^17^).

Studies on motor imagery have typically investigated the behavioral and brain responses to the mental rehearsal of a specific target movement^4,18,19^. However, movements in daily life are usually organized in motor sequences that comprise multiple, individual movements. In sports, for example, preparatory movements are not necessarily biomechanically advantageous but rather prepare the athlete for a subsequent, specific muscle activation, rhythm or timing. For instance, think of a basketball player may bounce the ball in a particular way or a particular number of times before a free throw. Such trained motor sequences support motor learning by cueing the corresponding target movement. Evidence suggests that the brain stores representations of entire motor sequences ^20^ and that individual motor segments affect the execution of the linked segments that follow ^21^. For example, letters are handwritten slightly differently depending on the preceding letter.

Motor adaptation refers to the process by which the motor system adjusts its output to compensate for changes in the environment or the body. When participants experience two randomly occurring perturbations during a reaching movement, e.g., in an interference force field task, they do not exhibit adaptation, given the perturbations’ randomness. Yet, if each perturbation is associated with a unique prior movement, participants can successfully adapt their movements ^22^. This kind of learning relies on associations between the kinematics of the prior movement and the adjustments required for the target movement, that is, the perturbed reach. The smaller the variability of the associated prior movement, the stronger the participants adapt^23^. Thus, the execution of a perturbed target movement benefits from a specific, regularly preceding movement in the sequence. Given the importance of sequence storage for learning and execution of linked overt movements, we asked whether a similar mechanism is at play when prior movements of a sequence are imagined rather than executed. Previous studies proposed that overt movement and motor imagery are functionally equivalent^18,7,24,25^. This suggests that overt segments in a motor sequence can be substituted with motor imagery without altering the overall outcome of the sequence. To test this, we integrated either executed or imagined prior movements with a perturbed overt target reach within a single sequence in an interference force field task. Participants successfully adapted their reaching movements to the interfering perturbations both when the linked, prior movement was executed or imagined. However, the adaptation was reduced for the imagined group. Moreover, power changes in the alpha and beta band during an independent basic motor imagery task, such as imagined fist clenching, were correlated with adaptation performance in the interference force field reaching task.

## 2 Results

Participants made reaching movements to different targets in an Exoskeleton Lab (Kinarm, Kingston Ontario). This device allows administration of precise perturbations of arm movements (see Fig. 1A). Participants were randomly assigned to one of three different groups (see Fig. 1B). Participants of the control group made a single reach from a middle to a final target. The position of an additional cue target indicated the possible perturbation direction during each trial. Participants of the active group performed a movement sequence that started with a reach from the cue’s position to the middle target, followed by a reach from the middle to the final target. Participants of the motor imagery (MI) group initially positioned their hand at the middle target but were instructed to perform kinesthetic imagery of a movement from the cue to the middle target, and to subsequently execute the reach from the middle to the final target. In this way they performed a hybrid sequence that encompassed both an imagined and an executed segment. The start signal for each (imagined or overt) reaching movement was a color change of the cue and middle target, respectively (see Fig. 1C). The reaching task was performed in blocks: 6 baseline, 50 adaptation, and 4 washout (see Fig. 1D). Each block comprised 18 trials. In the baseline phase, reaches were not perturbed. During the adaptation phase, a velocity-dependent, curl force field perturbed the reach between the middle and final targets. The direction of the force field was associated with the location of the cue in relation to the final target and therefore the direction of the (imagined) prior movement in the active and MI groups (see Fig. 1E). No group was informed of this relation and, according to post-task questioning, participants were not explicitly aware of the association between the force field direction and the cues’ location. In the washout phase, reaches were no longer perturbed. This allowed measurement of after-effects induced by the adaptation procedure.

**Figure 1:**
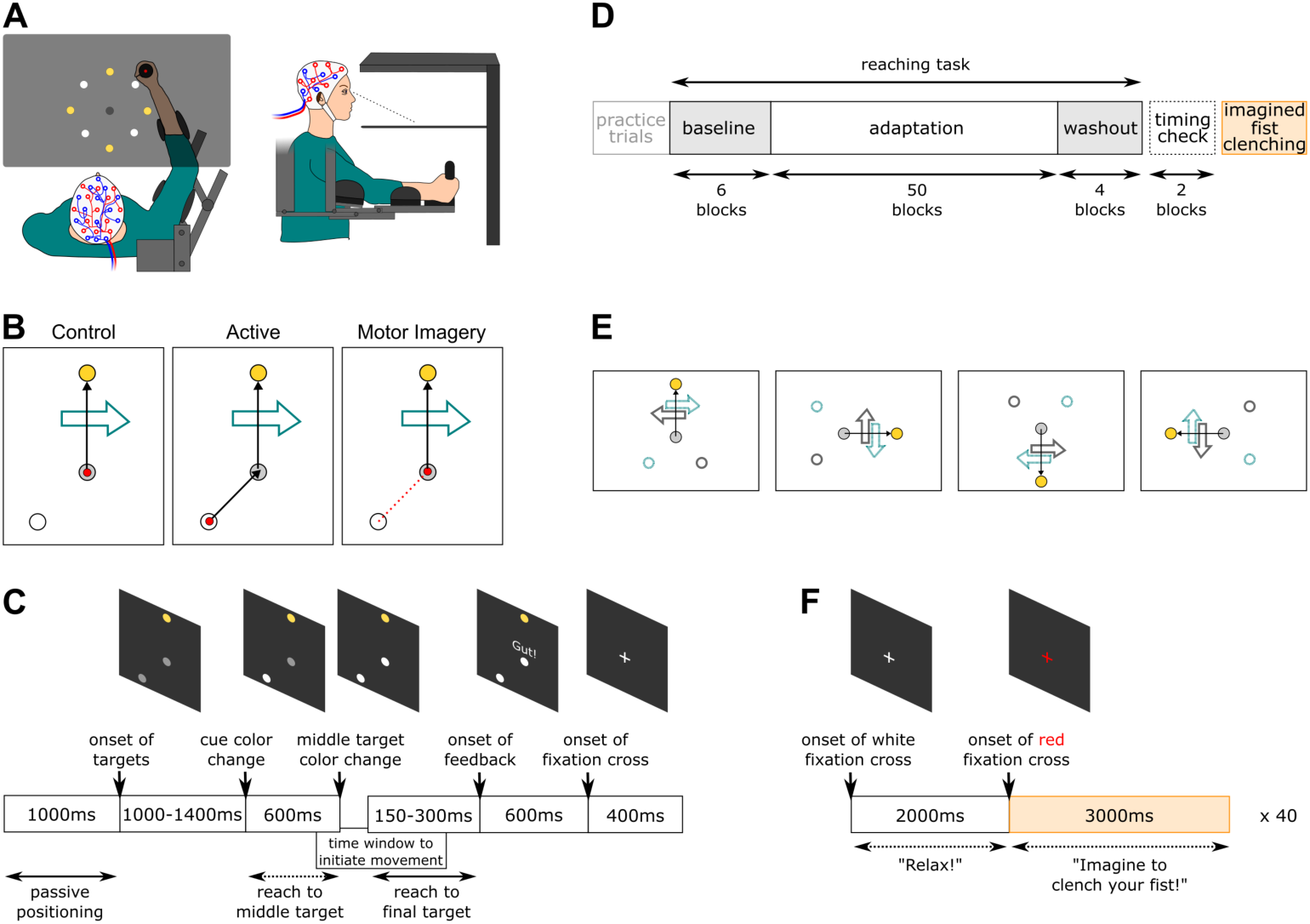
Experimental design and protocol. A) Experimental setup. The Kinarm Exoskeleton Robot Lab features a handle positioned underneath a horizontal screen on which targets are presented such that they appear to be located in the same horizontal plane as the hand. The middle target location is shown in black, the possible final target locations in yellow, and the corresponding cue positions in white. Current hand position was continually displayed as a red dot. In each trial, the middle target, one of the four possible final targets and a perturbation cue indicative of perturbation direction – always a dot positioned 45° clockwise or anti-clockwise of the final target, respectively, see panel E – were displayed. The screen/mirror is displayed transparent here for illustration; however, in the experiment participants were not able to see their arms. Left: Bird’s-eye view. Right: Side view. B) Exemplary trial of the reaching task separately for each experimental group. White dot - cue, grey dot - middle target, yellow dot - final target, red dot - hand position at the beginning of a trial, black arrow - instructed reaching path during the trial, red arrow - imaginary reaching path, big teal arrow - exemplary force field direction. C) Reaching task trial sequence. The prior movement, that is, the reach from cue to middle target was only overtly performed by the active group. The imagery group only imagined the prior movement while the hand was already positioned at the middle target, at which the reach to the final target would start. The control group did not perform or imagine a prior movement at all. Feedback about the movement time between middle and final target was given after every trial to encourage a similar reaching speed across participants. D) Experimental phases. The timing check task consisted of active reaches regardless of group membership. The imagined fist clenching task was only performed by participants in the MI group. E) All cue/final target combinations with big arrows representing force field direction depending on the cue’s location. The relationship between cue location and force field direction was reversed for half of the participants. White dot with teal/grey border - cue, grey dot - middle target, yellow dot - final target, black arrow - desired reaching path, big teal/grey arrow - force field direction. F) Imagined fist clenching task sequence. Participants were instructed to relax or to perform kinesthetic motor imagery of clenching their fist around the Kinarm’s handle when the fixation cross turned white or red, respectively. A, B, C, D, and E were adapted from ^26^.

### Kinematic results

Reaching trajectories were curved in the direction of the force fields at the beginning of the adaptation phase (see Fig. 2A). The active group adjusted their reaches to counteract the forces by the end of this phase, while the control group’s reaches remained strongly perturbed. This result pattern replicates previous findings and confirms that participants can adapt to multiple environmental perturbations when these are disambiguated by prior segments in a longer movement sequence (e.g., ^22^). The key finding is that, like the active group, the MI group exhibited adaptation. However, the adaptation was weaker, with significant curvature of reaches remaining even after all 50 blocks of the adaptation phase. Thus, adaptation of the imagery group appeared to be intermediate between the active and control groups. Similarly, after-effects in the washout phase, manifested by deflections of reaches in the opposite direction of the previously experienced force fields, were strong in the active group, less strong for MI, and virtually absent in the controls.

**Figure 2:**
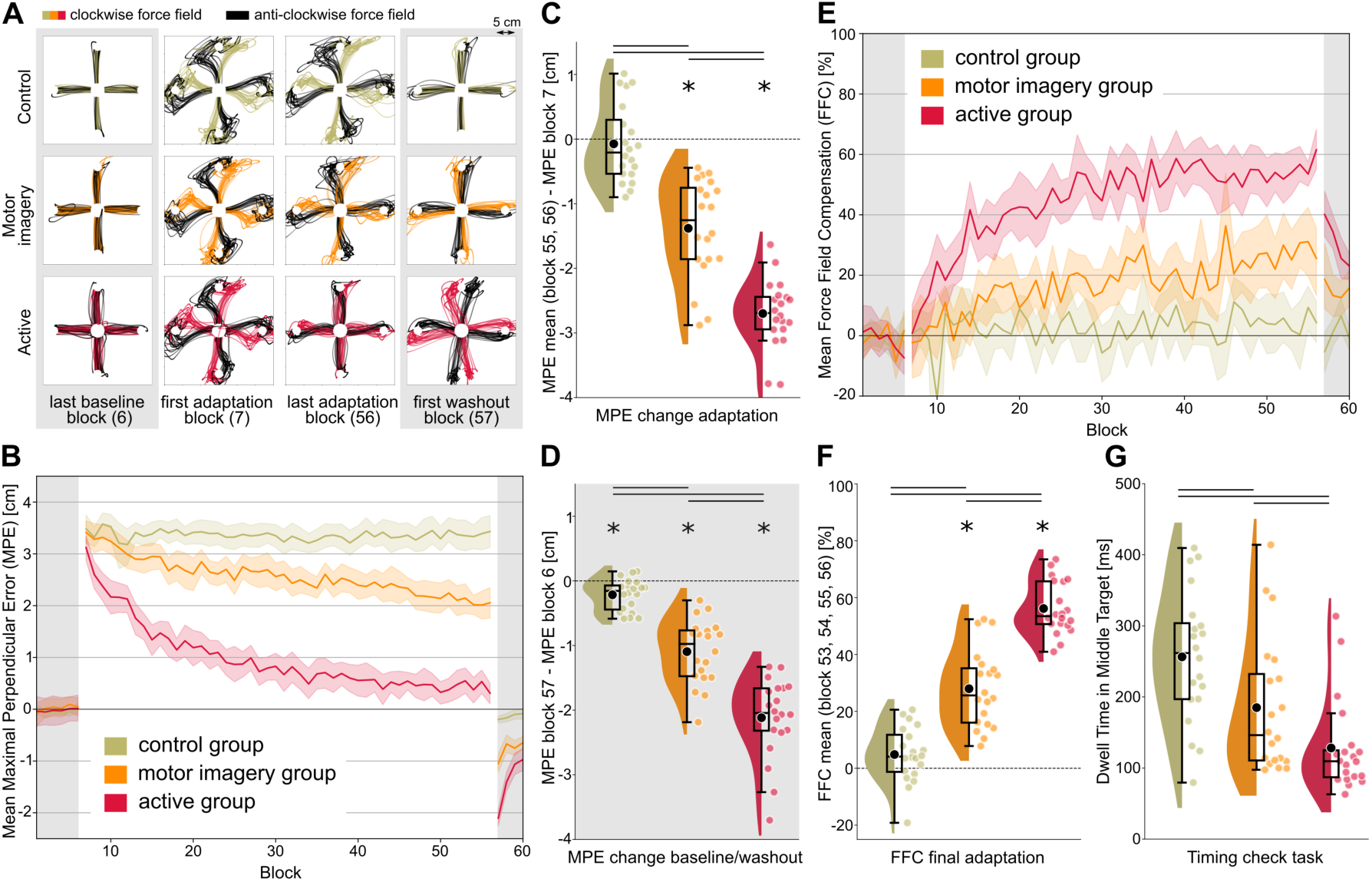
Behavioral results. Data of the Control and Active groups were also reported in ^26^. A) Reaching trajectories. Single trial trajectories from middle to final targets of all participants (N = 20 in each group) in specific blocks of the experiment. For each final target/cue position combination, one reaching trajectory per participant per block is shown. Force fields were only present in adaptation blocks. B) Averaged maximal perpendicular error (MPE) over trials and participants for each block. Force fields were present from block 7 to block 56 (adaptation blocks = white background). Error bands indicate 95% CIs. C) Change of MPE from beginning to end of adaptation. Each colored dot depicts the difference between average performance in the first and last two adaptation blocks of one participant. Black dots mark the respective group averages. Lines denote significant differences between groups *p <* .05. Stars mark significant within-group effects *p <* .05. D) Change of MPE from baseline to washout. The differences between the first washout and the last baseline block are displayed per participant and as a group average. E) Averaged FFC over trials and participants for each block. F) Average FFC in the last 4 adaptation blocks. G) Comparison of median dwell time in the middle target in the timing check task. Each colored dot depicts the median dwell time of each participant.

To systematically quantify the extent of adaptation we calculated the maximal signed deviation from a straight line between the middle and final targets for each reach. This Maximal Perpendicular Error (MPE) was then averaged over trials within each block (see Fig. 2B). We compared the MPE at the beginning and end of the adaptation phase within and across groups. Paired t-tests within each group revealed that MPE change during the adaptation phase was significant in the active (*t* (19) = 22.87, *p* = 1.3e-14; all reported p-values for t-tests are corrected for multiple comparisons) and MI groups (*t* (19) = 8.01, *p* = 6.5e-07) but not in the control group (*t* (19) = 0.54, *p* = 0.59). Active and MI participants exhibited significantly reduced error at the end of the adaptation phase. Still, t-tests between groups were significant for all three pair-wise comparisons (*t* (18) = −14.60, *p_active/control_* = 2.2e-16; *t* (18) = −6.32, *p_active/MI_* = 6.3e-07; *t* (18) = −5.96, *p_MI/control_* = 1.3e-06). This confirms that MPE change was strongest in the active group, but that MI participants also performed better than control group participants (see Fig. 2C). Error reduction, as measured by MPE change in the adaptation phase, could be due to compensatory mechanisms specific to the force fields (i.e., real motor adaptation) or by simply stiffening the reaching limb. To eliminate the possibility of such a generic and non-directional strategy, we compared the difference in MPE between the end of baseline and beginning of washout within and across groups. Participants who adapted to the force fields should have exhibited after-effects once the force field was removed (e.g., ^27^). Conversely, if participants had merely adopted a stiffening strategy, no deviation from straight reaching should be observed when no force field is present. Indeed, all paired t-tests within groups were significant (*t* (19) = 15.21, *p_active_* = 2.1e-11; *t* (19) = 9.82, *p_MI_* = 2.8e-08; *t* (19) = 3.97, *p_control_*= 0.0008), indicating that after-effects were present, albeit to a different extent. Participants in the active group showed greater after-effects than participants in the MI (*t* (18) = −5.73, *p* = 2.7e-06) and control group (*t* (18) = −12.74, *p* = 1.6e-14); and participants in the MI group showed greater after-effects than participants in the control group (*t* (18) = −7.10, *p* = 5.3e-08; see Fig. 2D). Contrary to the comparison of performance at the beginning vs. end of adaptation, this result suggests that even control participants were able to benefit from the visual cue to counteract the interfering force fields to a small extent. Imagining prior movements, however, allowed a much stronger adaptation, and performing overt prior movements resulted in the strongest effects.

Another measure that is often reported in adaptation experiments refers to the predictive compensation participants exhibit after experiencing force fields (e.g., ^22^). In our reaching tasks, two trials of each block were clamp trials, in which the exoskeleton robot restricted reaches to a straight line from the middle to the final target. We measured the force the robot needed to apply to keep participants on the straight trajectory. If a participant had learned to counteract a force field, then the force the exoskeleton robot must apply in clamp trials should be of equal magnitude. Accordingly, a value of 100% force field compensation (FFC) would indicate that a participant learned to perfectly counteract the previously experienced force fields. We observed the same adaptation pattern of averaged FFC per group over blocks as with the MPE (see Fig. 2E, F, and supplementary Results 1). Taken together, imagining a prior movement, associated with force field direction, allowed motor adaption, albeit less robust than explicitly making the prior movement. The same perceptual information, without a prior movement (imagined or executed) led to minimal adaption. Motor imagery is thus an effective contextual cue for force field-specific learning and can substitute for an active prior movement to some extent.

### Timing check task

Next, we asked whether a hybrid practiced sequence would enhance performance of the whole overt sequence production. To approach this question, all participants performed two active reaches in a timing check task. Instructions were identical to those for the active group in the main reaching task (see Methods). We compared, between groups, the median reaction times and dwell times in the middle target. A permutation test was used due to skewed data.

All groups initiated their reaches to the middle target similarly fast (all group comparisons for reaction time medians *p >* 0.16; *Md* _active_ = 379.75 ms; *Md* _MI_ = 396.25 ms; *Md* _control_ = 373.25 ms). However, we reasoned that faster transitions between the segments should be apparent after a sequence is learned ^28^. Accordingly, we analyzed how long participants dwelled at the middle target in the timing check task. Participants in the active group spent less time at the middle target than the MI (*p* = 0.0321) and control groups (*p* = 5.2e-05). This is not surprising, given that the timing check task was identical to the main reaching task for active participants and, thus, they had ample training. Critically, however, the MI group also dwelled for a shorter time at the middle target than the control group (*p* = 0.0403). Even though perceptual information had been identical for the two groups in the main reaching task, MI participants were faster to connect the two reaches in the timing check task. This indicates that participants in the MI group used the hybrid sequence as training for the full execution of the same sequence, implying that they formed a representation of two linked movements in the reaching task, even though they only imagined the first segment. The imaginary training, thus, allowed the MI group to connect the two overt movements in the timing check task faster than the control group.

### Neural oscillatory changes during motor imagery

Next, our objective was to examine the potential association between neural markers of motor imagery proficiency and adaptation performance. For this, we recorded electroencephalography (EEG) with 60 electrodes and electromyography (EMG) of right arm muscles during the entire experiment. To measure neural activity during motor imagery, without interfering signals from motor planning for an upcoming reach, participants of the MI group additionally performed the imagined fist clenching task (see Fig. 1D). At the start of this task, a white fixation cross was displayed. After 2 seconds, the cross turned red, signaling participants to imagine tightly gripping the handle with their right hand for 3 seconds, until the cross turned white again (see Fig. 1F). This sequence of events was repeated 40 times.

We focused on changes in the power of oscillations in alpha and beta frequency bands, which were previously shown to be modulated by motor imagery ^5,29,30^. Therefore, we analyzed the power change during motor imagery starting at 0 s relative to the baseline activity, which was calculated in the interval −0.75 s to −0.25 s. In line with previous studies, we observed a lateralized event-related desynchronization (ERD), in alpha and beta bands, in electrodes over the contralateral sensorimotor cortex after the start of the mental imagery averaged over participants (see Fig. 3; for more details see supplementary Results 2).

**Figure 3:**
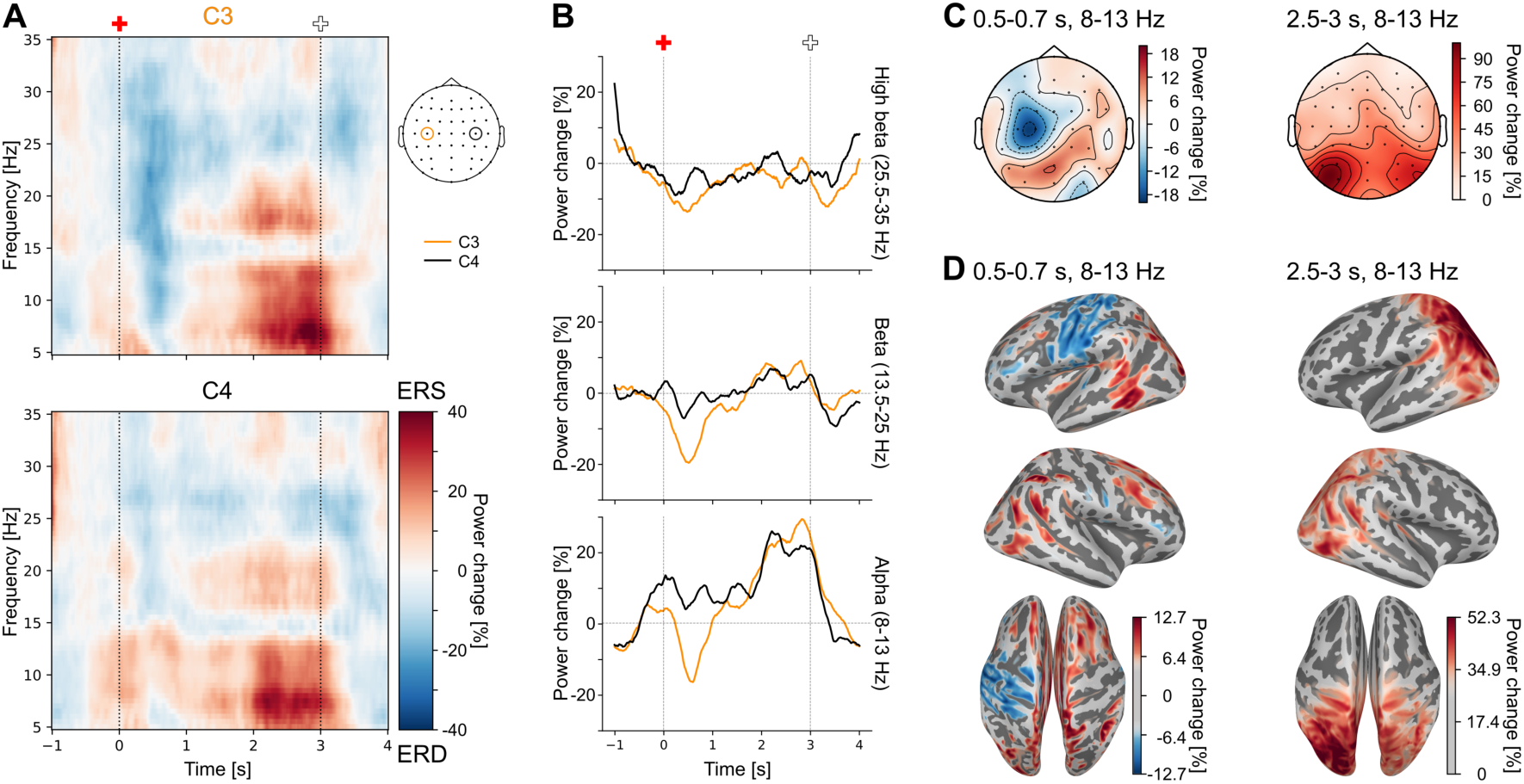
EEG data imagined fist clenching task. A) Time-frequency representation of grand averaged (N = 16) data at electrode standard positions C3 and C4. Color represents power change in percent relative to the baseline window from −0.75 to −0.25s. Red cross - start of motor imagery, white cross - start of relaxation. B) Power change in percent over time relative to baseline activity in 3 frequency ranges averaged across participants and respective frequencies. Orange line - C3 electrode; black line - C4 electrode. C) Distribution of averaged alpha activity across the scalp in specified time windows (left - ERD; right - ERS), averaged across participants. D) Source reconstruction of averaged alpha activity in chosen time windows, averaged across participants. The strongest 50% of values are shown in color.

In the more complex reaching task, we did not observe this typical lateralized motor imagery pattern (see supplementary Fig. 2), likely reflecting an overlap of neural activity resulting from imagined, planned, and executed movements.

### EEG of imagined fist clenching task predicts motor adaptation

To assess how neural data in the imagined fist clenching task related to adaptation performance in the reaching task, we first linearly combined our 3 behavioral adaptation measures into an overall measure of error reduction to maximize robustness and reliability. This included the change of MPE from the beginning to end of adaptation, the change of MPE between baseline and washout, and the FFC at the end of adaptation. Next, we tested the correlation of this change of error with each time-frequency bin in all channels across participants and performed a cluster-based permutation test (initial grouping threshold = 0.01, 1000 permutations) to investigate the relationship between oscillatory brain activity during the imagined fist clenching task and adaptation performance.

Change of error was negatively correlated with power change (*p*_cluster_ = 0.023). This correlation was driven by power changes from approximately 2 to 3.7 s, which, thus, included a time interval after the termination of motor imagery at 3 s. The cluster comprised both alpha and beta frequency ranges and was most prominent in channels C3 and F3 (see Fig. 4 A & B; for topoplots displaying unthresholded data see supplementary Fig. 3A). The single strongest correlations of all single time-frequency bins in C3 and F3 were *r* = −0.891 (at 34.5 Hz, 3.15 s) and *r* = −0.84 (at 8.5 Hz, 3 s), respectively. Negative correlations indicate that bigger oscillatory power increases were associated with more negative changes of error (i.e., stronger performance improvement) in the respective time-frequency-channel bins belonging to the significant cluster. Co-aligning the averaged time-frequency representation (for example C3, see Fig. 3A) with the significant cluster (see Fig. 4A) indicated that a stronger event-related synchronization (ERS) was related to a bigger performance improvement observed during adaptation. We observed similar correlation patterns (lateralized and strong correlations at C3) when we assessed the relationship between the neural data and individual behavioral measures (see supplementary Fig. 4).

**Figure 4:**
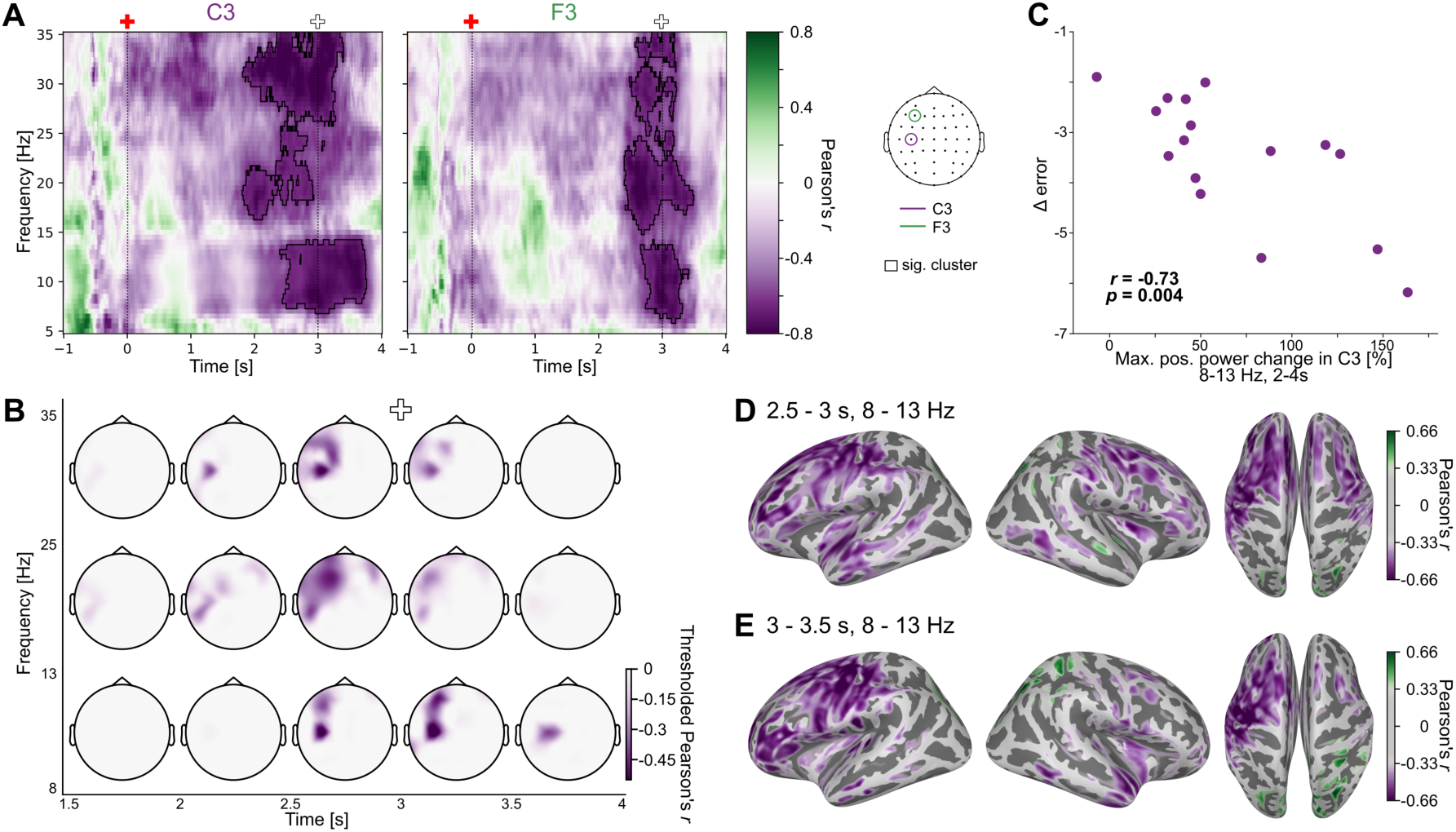
Correlation analysis results. A) Pearson’s correlation of change of error in the reaching task with power change in the imagined fist clenching task across participants in C3 and F3. The pixel color indicates the respective correlation value at that location. A more negative change of error denotes better adaptation performance. Red cross - start of motor imagery, white cross - start of relaxation. The significant cluster is outlined in black. B) Topoplots of different time windows and frequency ranges displaying the distribution of thresholded correlation values. Only the significant cluster is shown. Correlation values are averaged in the specified time and frequency ranges. C) Relationship between change of error in the reaching task and the maximal positive power change value (ERS) in the imagined fist clenching task across participants. Each point depicts a participant. The maximal positive power change value was taken for each participant in the alpha frequency range between 2 and 4 s. D)&E) Pearson’s correlation of change of error in the reaching task with power change in the imagined fist clenching task - averaged over specified time and frequency range - across participants in source space. The strongest 50% of values are shown in color in D and the color bar is kept identical for E.

Because different participants may exhibit their peak ERD and ERS responses at different times in our analyzed time window, we tested whether individual alpha peak ERD or ERS values in channel C3 correlated with error reduction across participants. We chose the alpha frequency range for our main analysis because here we observed the largest power modulation (see Fig. 3C; for beta see supplementary table 1). The peak ERD did not correlate with the change of error across participants (*r* (14) = −0.116, *p* = 0.67). The peak ERS, however, strongly correlated with change of error (*r* (14) = −0.73, *p* = 0.004, see Fig. 3C). Similarly, the overall measure of oscillatory responses based on the peak-to-peak difference of ERD and ERS in each participant correlated with change of error (*r* (14) = −0.722, *p* = 0.003). Thus, stronger ERS in the imagined fist clenching task and a stronger overall modulation of the alpha band power were both predictive of motor adaptation performance in the reaching task. Because the sample size for the correlation analyses was relatively small, we undertook additional analyses to demonstrate that our findings are robust over different analysis pipelines and not dependent on single participants (see supplementary Table 1).

### Source reconstruction

To more precisely identify brain sources, we visualized the correlations of averaged power values in specified time/frequency windows with change of error in source space. In the ERS time window (2.5 - 3s) the strongest negative correlations were mainly found in the left postcentral and superior-frontal regions (see Fig. 4D). In the time window right after completion of the imagined fist clenching (3 - 3.5s) the strongest negative correlations were observed in the left post- and pre-central regions (see Fig. 4E; see supplementary Fig. 2B, C, D for correlation’s strength over additional time/frequency ranges).

### No evidence that EEG of reaching task or subjective rating of motor imagery predicts motor adaptation

In a next step, we searched for a relationship between oscillatory activity during the imagined movement in the reaching task and motor adaptation performance. However, we did not find a significant correlation across participants when performing the same analysis steps as we did with the imagined fist clenching EEG data (see supplementary Results 4). This is likely due to motor imagery and planning processes overlapping in the reaching task and therefore obscure the relationship between oscillatory dynamics and adaptation performance. In addition, we did not find a relationship between perceived vividness in motor imagery and motor performance or neural data (see supplementary Results 5). This suggests, that motor imagery proficiency might not be reliably accessible by subjective ratings.

## 3 Discussion

Motor imagery is a widely used tool to facilitate motor control and learning. Typically, the target movement is imagined to improve exactly the same movement. In this study, we show that motor imagery can be a useful technique beyond improving just the imagined movement. We present kinematic evidence that an imagined, prior segment of a sequence aids adaptation to perturbations and leads to faster transition between segments of the actual, full execution of the sequence. Our findings imply that hybrid sequences, that is, sequences comprising both overt and imagined segments, are stored as a single, holistic unit, akin to fully executed motor sequences ^20^. At the neural level, the linkage effects we observed were related to well-established motor mechanisms, though the precise role of this region still needs to be clarified.

Previous work has already shown that imaginary movements that occurred after (rather than, here, before) the force field-affected reach improved adaptation ^31^. However, when an imagined segment is instructed to occur last, participants may have planned the full sequence only then to abort its execution just before the final, imagery element. Consequently, they may have associated a full-sequence motor plan with the expected force field ^32^. Our paradigm excluded this alternative strategy, thereby requiring genuine motor imagery generation. Our findings, including improved adaptation and reduced dwell-time between segments, demonstrate that motor imagery can have benefits beyond just benefiting target movement performance itself. By incorporating motor imagery in a partly executed motor sequence, training of this hybrid sequence can, on the one hand, facilitate selection of the desired motor response. On the other hand, linking of well-trained hybrid sequences can also transfer to overt sequence production.

Our results thus support the functional equivalence model, which postulates that motor imagery involves generating a complete motor plan that is merely inhibited from use at the execution stage ^24^. According to this model, neural representations of imagined and executed movements largely overlap, offering a potential mechanism for how imagining one sequence segment can affect performance and learning of linked elements. Here, the motor representation of a well-trained hybrid sequence provided the information required to counteract the forces that arise from dynamics that would be experienced later in the sequence. This is in line with motor imagery activating forward models to predict the associated, hypothetical sensory outcomes ^33^.

Beyond the evident difference in the physical execution and sensory feedback of the movement, overt and imagined movements are clearly distinct. For example, increased activity in one brain area during motor imagery does not necessarily imply increased activity during movement, or vice versa ^4,34^. In addition, behavioral differences after motor imagery practice and motor execution have been reported before ^35,36^. We observed lower adaptation performance for participants performing motor imagery compared to those actively performing the whole reaching sequence. We suggest this is the case because of the weaker neural activation during motor imagery as compared to overt movement^5^. Weaker activation is likely to have a less pronounced effect on the following movement execution and therefore on learning of the hybrid sequence. A recent ultra-high-resolution 7T study provided compelling evidence that motor imagery evokes responses in the superficial layers of the primary motor cortex (M1), whereas overt movement evokes responses in both superficial and deeper layers ^37^. This finding is in line with current concepts of layer-specific organization of M1. Superficial layers receive somatosensory and premotor input, whereas cortico-spinal output, needed for actual movement, is primarily derived from deep layers ^38^. Interestingly, in rats, the activation of deep layers in M1 seems to be critical for successful motor learning ^39^. Speculatively, then, the difference in adaptation performance we observed between participants performing an overt as compared to an imagined prior movement may result from varying activation patterns in the neural layers of M1, consistent with the strong, localized effect of imagery over the sensorimotor cortex in our EEG data. Whereas engagement of only superficial layers of human M1 may principally be sufficient for motor adaptation to occur (as in the motor imagery group), evoking responses from deep layers (when performing an overt movement) may further support adaptation.

We hypothesized that individuals with higher proficiency in motor imagery would experience greater benefits from imagining prior movements compared to those with lower proficiency. Stronger ERS in the imagined fist clenching task in both the alpha and beta bands was associated with a greater reduction of movement error and, therefore, stronger adaptation performance in the reaching task. Beta ERS following real or imagined movement in the sensorimotor region reflects active inhibition ^40,41^. In addition, alpha ERS signifies top-down inhibitory control mechanisms when subjects suppress or control the execution of a motor response ^42^. Thus, consistent with the functional equivalence model, better overall motor imagery ability may rely on a stronger inhibition response in the imagined fist clenching task. This, in turn, might lead to better linkage of imagined and real movement and therefore better adaptation performance in the reaching task.

Correlations between the power of oscillations in the imagined fist clenching task and motor adaptation were spatially widespread. A network involving contralateral sensorimotor and prefrontal regions showed strong correlations in both the alpha and beta bands. Activity in the dorsolateral prefrontal cortex (DLPFC) has been consistently reported during motor imagery ^4^ and is likely related to higher-level control processes. DLPFC activity has been related to movement inhibition ^43^, fitting with the functional equivalence model’s requirement to forego execution of planned movements. Others have attributed DLPFC activity to adaptive cognitive control ^44^, in line with the motor-cognitive model of motor imagery, which suggests that imagining a movement uses more executive resources than actually performing the movement, due to the lack of sensory feedback ^45^. Moreover, the motor simulation theory postulates that higher cognitive systems interact with and supervise the motor simulation process ^25^. The lack of sensory feedback may necessitate the engagement of higher-level control processes during imagined execution. The ERS we observed in the fist clenching task may, thus, reflect sufficient engagement of executive control during imagery to simulate the movement. As motor adaptation also relies on executive functions ^46^, superior executive function may be associated with both stronger ERS in the imagined fist clenching task and better adaptation performance in the reaching task. Finally, successful motor adaptation requires the somatosensory cortex ^47^. The ability to form and update motor memories might be reflected by both the adaptation performance in the reaching task and by synchronization of oscillatory activity in the somatosensory cortex after an imagined fist clench, contributing to the observed correlation pattern.

Our findings, including improved adaptation and reduced dwell-time between segments, demonstrate that motor imagery can have benefits beyond just benefiting target movement performance itself. By incorporating motor imagery in a partly executed motor sequence, training of this hybrid sequence can, on the one hand, facilitate selection of the desired motor response. On the other hand, linking of well-trained hybrid sequences can also transfer to overt sequence production.

Our results open exciting possibilities for sports training and rehabilitation. For example, in stroke recovery, gross motor skills, such as arm reaches, often recover before fine motor skills, such as finger movements ^48^. In this context, training a hybrid sequence consisting of, for example, an overt reach to an object and an imagined grasping movement (or vice versa), may facilitate the re-learning of the grasping movement through the well-researched effects of motor imagery of the target grasping movement (e.g., by stimulating the same neural pathways as actual movement and promoting brain plasticity). Moreover, with practice, the prior reaching movement can cue the specific (imagined) grasping movement and thereby further facilitate the learning process. Finally, the transition between reaching and grasping might be improved if both movements are overtly performed in sequence after hybrid training.

It will be crucial to assess which types of movements are suited for applied hybrid sequence interventions, and to determine what the best training approaches are. In the given example, patients might benefit more if they imagine (or execute) a specific prior reach trajectory linked to a specific (imagined) object grasp, or, alternatively, if they practice various sequences. Future research should also identify individual differences and prerequisites for effective interventions. In our study, subjective vividness of the imagined movement did not relate to performance improvement or to a neural correlate. This suggests that self-report might not sufficiently capture the aspects of motor imagery that relate to performance ^49,31^.

To summarize, our study demonstrates that hybrid sequences combining overt and imagined movements can significantly enhance adaptation to interfering force field perturbations. Accordingly, motor imagery can be leveraged to improve the performance of not only the imagined but also of linked overt movements. In addition, we have established a clear connection between motor adaptation ability and neural oscillatory dynamics observed during an unrelated motor imagery task. Notably, individuals who exhibited stronger oscillatory modulation during a basic motor imagery task showed greater adaptation improvement in overt movements that were linked to prior imagined movements. This finding underscores the importance of individual differences in motor imagery proficiency, suggesting that the effectiveness of imagery-based motor adaptation may be influenced by these neural dynamics.

## 4 Materials and Methods

### 4.1 Participants

A total of 65 volunteers aged 18-35 years participated in our study. We excluded 5 participants from our analysis, as explained below. Therefore, our final sample consisted of 60 participants (30 females, 30 males) with a mean age of 26.1 (*SD* = 4.6) years. All participants were right-handed and had normal or corrected-to-normal vision and no known neurological, perceptual, or motor impairments or disorders. The study received ethical approval from the local ethics committee at the University of Leipzig, and all participants provided informed written consent prior to the experiment.

### 4.2 Apparatus and Stimuli

#### Kinarm robot

The tasks were performed within a Kinarm Exoskeleton Lab (Kinarm, Kingston, Canada). This robotic device can reliably track arm movements in the horizontal 2D plane with a recording rate of 1000 Hz. Targets and individually calibrated hand positions are displayed via a mirror reflecting a monitor mounted above (see Fig. 1A).

#### EEG

We measured EEG with 60 passive electrodes (Brain Vision by Brain Products, Gilching, Germany) following the international 10-20 system. In addition, we measured electrooculogram (EOG) with 4 electrodes, electrocardiogram (ECG) with 2 bipolar electrodes, and EMG of the right arm and shoulder muscles (brachioradialis, triceps lateral head, pectoralis major, posterior deltoid) with 8 bipolar electrodes. The sampling rate for all recorded biological signals was 2500 Hz. All electrodes were attached to the participants before they entered the Kinarm robot.

### 4.3 Tasks and Procedure

Participants were randomly assigned to the control, active, or motor imagery (MI) group (see Fig. 1B). Each participant performed a reaching task that was slightly different depending on the group. The subsequent timing check task was the same for everyone. Participants in the MI group additionally performed the imagined fist clenching task (see Fig, 1D & F). Lastly, participants were asked to fill out a short questionnaire.

#### Reaching task

Participants performed right-arm reaches towards targets in the Kinarm robot. Participants were given a brief introduction to the task before starting the experiment. They then completed a total of 60 blocks, consisting of 6 baseline, 50 adaptation, and 4 washout blocks (see Fig. 1D). Each block included 16 normal and 2 clamp trials. During normal trials, in the adaptation phase, force fields perturbed the reaching movement between the middle and final targets. The robot’s perturbations in the adaptation phase appeared random to the participants included in this study.

In total, participants performed a minimum of 1080 trials. Short breaks were taken approximately every 200 trials, and there was also a 5-minute break at the halfway point. Example videos of the reaching task of each group can be found on OSF (https://osf.io/swgd9).

During each reaching trial, three targets each with a diameter of 1.25 cm were presented: the cue, middle, and final target. The location of the middle target was adjusted based on the individual’s hand position when the elbow was bent at a 90° angle and the shoulder angle was at 60°. There were four potential positions for the final target, located either 12 cm to the right, left, up, or down from the middle target. For each final target position, there were two possible cue positions (see Fig 1E). The distance between the cue and the middle target was 10 cm. The current hand position was indicated by a red cursor with a diameter of 0.5 cm. The cue and middle target were gray and changed to white during the trial, and the final target was yellow. The targets’ number, color, size, timing, and between-target angles were determined based on previous research (e.g., ^22^ and the same as in ^26^).

Depending on the group, a trial consisted of either one or two active reaches. In every trial, all experimental groups performed the same final reach, which involved moving the right hand from the middle to the final target. A velocity-dependent curl field was sometimes present between the middle and final target causing systematic perturbations to the arm during those reaches. The force field began after the right hand was more than 2 cm away from the middle target’s midpoint and remained present until the final target was reached. The location of the cue, relative to the final target, determined the direction of the force field. To account for any kinematic or biomechanic advantages, half of the participants learned that a positive angle between the cue and final target corresponded to a clockwise (CW) force field, while the other half learned that it corresponded to a counterclockwise (CCW) force field. This association between the angle’s sign and the direction of the force field remained constant for each participant throughout the experiment. The experienced forces *F* were perpendicular to movement direction and varied based on reaching velocity:

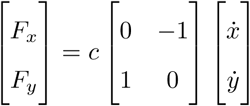

Depending on the location of the cue, the constant c was set to −13 or 13 Ns/m^50^. There was never a force field between the cue and the middle target. In randomly selected trials of the reaching task, the Kinarm robot enforced straight movements between the middle and left final target by using a force channel. We refer to these trials as clamp trials because the force channel walls restricted participants’ movements to a straight path. If participants adapted to the previously experienced force fields, they would anticipate the direction and strength of the fields and counteract the perturbations. In clamp trials, the Kinarm robot measured any compensatory forces applied by the participants against the channel walls. This allowed us to assess any potential feedforward learning. Due to technical limitations, the force channel walls were sometimes not strong enough. As a result, participants were able to slightly deviate from the straight line, and in some cases, they even broke through the virtual wall and moved off the intended trajectory. These trials were excluded from our analysis. Performance of the last four adaptation blocks was, thus, based on 407 instead of 480 total trials.

Each group consisted of 20 participants (10 females, 10 males). The motor imagery group had a mean age of 25.75 (*SD* = 4.46) years. During each trial in the motor imagery group, a white fixation cross appeared on a black background, and the robot moved the participants’ right arm towards the middle target (see Fig. 1C). This passive positioning lasted for 1000 ms. Following this, all targets (cue, middle, and final) and the hand position cursor were displayed. After a random period of time ranging from 1000 to 1400 ms, the color of the cue changed from gray to white. This was the signal for the motor imagery group to start imagining moving their hand from the cue position to the middle target. Participants were instructed to imagine the movement, or more specifically, to try to “feel” the movement and how this feeling differed depending on the cue’s location. To make this reaching imagery easier, participants executed the actual reach from the cue to the middle target in the practice trials, before the start of the reaching task. 600 ms after the cue color change, the color of the middle target changed from gray to white, indicating that participants should have concluded their imaginary reach and should start to actually move to the final target. Once they reached the final target, feedback on their movement speed was displayed just above the middle target. If the time to move from 2 cm away from the middle to the final target fell between 150 and 300 ms, the feedback (in German) was “good”. Otherwise, if it was faster or slower than this range, the feedback was “too fast” or “too slow” accordingly. The feedback was shown for 600 ms. Subsequently, a white fixation cross was displayed for 400 ms before the next passive positioning for the next trial began. When half of the trials in a block were completed, the inter-trial interval was extended to 4 s instead of 400 ms. If the cursor was not in the middle target when the color changes of the targets occurred, the trial was immediately aborted and repeated within the same block. Additionally, if participants left the middle target earlier than 100 ms before, or later than 500 ms after, the *go*-signal, the trial was considered unsuccessful and was repeated at a random position within the current block.

For two participants in the motor imagery group, the time between cue color change and middle target color change was 400 ms instead of 600 ms. However, their performance was comparable to the other participants in the group, with all three dependent variables (explained below) deviating by less than 1.4 standard deviations from the mean of the 600 ms group. Therefore, their data was included in the kinematic but not in the EEG analysis.

The timing, feedback, and repetition criteria were consistent across all groups. The control group was presented with the same trial procedure but was not told to do anything until the middle target changed color. In the active group, the participants’ hands were moved to the cue’s position at the start of the trial and they were instructed to make two subsequent reaches: from the cue to the middle target and from the middle target to the final target. The control and active groups have been previously described in greater detail in ^26^.

#### Timing check task

In the timing check task, all participants performed two reaches: from the cue to the middle and from the middle to the final target. The timing of color changes and abortion criteria were the same as in the reaching task. However, there were no clamp trials and no force fields. Another slight deviation was the position of the cue. In 33% of trials, the cue was moved to 75% of the original 10 cm distance from the cue to the middle target (7.5 cm) and in another 33% of trials, the cue was moved further away (13.3 cm), so that the original 10 cm distance equaled 75% of the longer distance. This is was done to introduce more variability and therefore difficulty in the task. There were two blocks with 24 trials each. Every unique cue/final target combination was presented three times, once for each tested distance between the cue and the middle target.

Due to the individual set-up and calibration, nine participants could not reach the cue’s position in the upper-left corner when it was moved further away (13.3cm) from the middle target. The non-executable trials were excluded and a block consisted of 22 trials for these participants. Two participants expressed the wish during the experiment to finish early and thus only executed one block instead of two.

#### Imagined fist clenching task

The imagined fist clenching task at the end of the experiment was only performed by participants in the MI group. Participants were still seated in the Kinarm robot. At the start of the task, the robot moved participants’ right hand to the former position of the middle target, where now a white fixation cross was displayed. After 2s the cross changed color from white to red and participants were instructed to imagine clenching their fist around the handle as hard as they could, as long as they saw the red cross. After 3s the cross changed back to white. The participants were instructed to remain in a relaxed state during the presentation of the white cross. This sequence was repeated 40 times.

We introduced the imagined fist clenching task after a break in data collection due to COVID-19 pandemic restrictions. For this reason, the first three MI participants did not perform the task. Another participant had to be excluded because they contracted their muscles during the 3 motor imagery seconds (captured by EMG, see below). The sample of the imagined fist clenching task consisted therefore of 16 participants.

#### Questionnaire

At the end of the experiment, participants filled out a questionnaire. One objective was to find out if participants had noticed any specific pattern between the force field direction and the cue position and whether they were explicitly aware of it. As a result, five participants (three from the MI group, two from the control group), who correctly identified this association, were excluded from the analysis and additional data from five new participants were acquired to replace them.

A second objective was to obtain a subjective rating from participants in the motor imagery group about their perceived ease/difficulty of imagining the movement in the reaching task. For this, we adapted the questions from the widely utilized motor imagery questionnaire (MIQ-RS, ^51^) to our task. Specifically, participants rated how well they thought they imagined the movement on average over the whole reaching task on a 7-point Likert Scale. The scale (in German) ranged from 1 (“extremely difficult to feel”) to 7 (“extremely easy to feel”) with a neutral option at 4 (“not difficult/not easy to feel (neutral)”).

### 4.4 Data analysis

#### Kinematic analysis

##### Preprocessing

The Kinarm device recorded angles of the elbow and shoulder joints at all time points (1000 Hz). We preprocessed the data in MATLAB (R2021a). A low-pass filter was applied to the data using a cutoff frequency of 10 Hz. We added hand velocity, acceleration, and commanded forces to the automatically recorded hand position. Commanded forces refer to the forces generated by the Kinarm device to ensure the participants’ hand remained within force channels during clamp trials. Our main analyses were conducted using Python (version 3.10) and relied on various libraries, including NumPy ^52^, Pandas ^53^, SciPy ^54^, scikit-learn ^55^, as well as Matplotlib ^56^ and Seaborn ^57^ for data visualization. We excluded all aborted trials in our analysis. Our preprocessing pipelines are described in Supplementary Methods 3.

##### Maximal perpendicular error (MPE)

One of the primary measures of interest was the Maximal Perpendicular Error (MPE), which quantified the maximal deviation in cm from a straight line connecting the middle and final targets of the right arm trajectory. A positive value indicated that participants exhibited a curved trajectory in the direction of the force field. The MPE was calculated from 2 cm past the middle target’s midpoint (toward the final target) and the final target’s endpoint. We excluded trials in relevant blocks in which it was evident that participants had chosen an incorrect target at the start (9 trials across all samples).

To assess how well each participant adapted to the force fields, we investigated two variables related to the MPE. First, we subtracted the average MPE of the first adaptation block from the average of the last two adaptation blocks (MPE change adaptation, see Fig. 2C). If the MPE change adaptation value was negative, it meant that the participant’s reaches became straighter (less perturbed by the force field) at the end of the adaptation phase compared to the beginning. A more negative value indicated a greater improvement in performance. Second, we calculated the difference between the average MPE of the last baseline and the first washout block (MPE change baseline/washout, see Fig. 2D). If this value was negative, it meant that the participant made consistently more curved reaches in the washout block in the opposite direction of the force field experienced before, compared to the baseline block. This would have been due to a residual counteracting effort that was applied earlier. Consequently, a more negative value indicated a larger force field after-effect.

We performed t-tests between groups and within groups, against zero, for both MPE change adaptation and MPE change baseline/washout. We applied the Bonferroni-Holm correction to adjust the p-values of each family of tests involving the same dependent variable (6 tests each).

##### Force field compensation (FFC)

A second outcome measure was force field compensation (FFC) in clamp trials. To determine FFC, we analyzed force data within a 150 ms time window centered on the time of peak velocity during the reach to the final target. Based on the movement’s velocity, we calculated the ideal force profile, which would have counteracted a present force field. We linearly regressed the measured force against the channel walls on the ideal force profile with the intercept forced to zero. We then defined FFC as the slope of the regression multiplied by 100%.

Next, we determined the average FFC for each participant in the final four blocks of the adaptation phase (referred to as FFC final adaptation, see fig. 2F). A final adaptation value of 100%, would mean that the participant made perfect adjustments to their reaches to counteract the force fields. We used FFC final adaptation to again perform t-tests within and between groups and adjusted with the Bonferroni-Holm correction (6 tests).

##### Timing check task

In the timing check task we defined reaction time as the time between the *go*-signal, indicated by the color switch of the cue, and the time when participants left the cue location. Dwell time, on the other hand, was defined as the time between entering and leaving the middle target. For each participant, we recorded the median reaction and median dwell time of all performed trials. We compared the two measures between groups with permutation tests. For this we randomly permuted the group labels and calculated the difference of medians one million times to create a null distribution for each comparison. The observed differences between groups were then compared to the respective null distributions to determine the p-values. Note that we did not correct for multiple comparisons here. This analysis was not preregistered and hence exploratory.

#### EEG analysis imagined fist clenching task

We performed our EEG data analyses using MNE Python ^58^ and custom scripts.

##### Preprocessing

The raw EEG data were downsampled to 250Hz (using the *resample* function of MNE Python, which applies a low-pass anti-aliasing FIR filter) and then re-referenced to the average reference. Next, the data were bandpass filtered between 0.5-45 Hz to remove unwanted frequency components using an 8-order Butterworth filter applied forward-backward to prevent phase distortions in data. Subsequently, a notch filter was applied to remove the 50 Hz power line noise with a frequency range of 50 Hz to 125 Hz, filtering the first and second harmonics. Noisy channels were excluded by visual inspection of data for each participant. The maximal number of excluded channels per participant was three.

Independent Component Analysis (ICA) was used to remove noisy components from data. For this, a copy of the data was filtered between 1 and 45 Hz to remove slow-frequency drifts ^59^. Independent components were estimated with MNE Python using the method *picard* and the fitting parameters set to obtain extended Infomax solutions. The algorithm included a principal component analysis (PCA) to whiten the data before performing ICA. The number of components was set to the number of available EEG channels minus one. Noisy ICA components were removed by visual inspection. Artifactual components included mostly eye movements and muscle activity. The remaining components were projected back to the sensor space. Finally, the continuous data were then segmented into epochs ranging from 1.5 seconds prior to the onset of motor imagery to 4.5 seconds afterward. Noisy epochs and epochs in which the EMG showed muscular activity in the right arm in the imagery compared to the relaxation phase were removed. One participant had to be removed entirely because they clearly contracted muscles in each motor imagery section. Finally, excluded bad channels were interpolated using MNE Python with *spline* method.

##### Time-frequency transformation

We computed the current source density (CSD) to estimate spatially specific neuronal activity. To analyze changes in neural oscillations over time and across various frequency bands, we used MNE Python to compute the time-frequency representation (TFR) of the data. A multitaper approach was applied to calculate the power of neural activity for each frequency bin and time point^60^. The frequency range for the TFR was set between 5 and 35 Hz, with a step size of 0.5 Hz. The number of cycles for each frequency bin *f* was set to *f /*2. Within each participant, we calculated the average power across epochs in each frequency-time bin. To reduce memory usage we used a decimation factor of 2 after time-frequency decomposition. We cropped the resulting TFR data from −1 to 4 s.

We performed a baseline correction by expressing the TFR data as a percentage change relative to the mean baseline activity from −0.75 to −0.25 s in relation to the start of motor imagery. This way, we ensured that the subsequent analyses focused on changes in neural activity relative to a time interval in the relaxation phase prior to the motor imagery onset.

##### Source reconstruction

To obtain more precise information about the spatial location of neural sources, we mapped our data to source space. A forward solution was calculated based on *fsaverage* standard head model, the three-layer boundary element model (BEM), and the dipole grid with ico5 spacing accompanied by MNE Python ^61^. The resulting source space consisted of 10242 dipoles per hemisphere. We computed the Minimum Norm Estimate (MNE) ^62^ inverse solution using MNE Python, with dipole orientations normal to the cortical surface and the regularization parameter equal to 0.05. Next, we performed the same time-frequency analysis on the source estimates in source space as on the sensor data (see time-frequency transformation). We used a frequency range of 8-13Hz for the alpha band, and 14-25Hz for the beta band, with a step size of 1 Hz. We used the same baseline correction approach as in the sensor data analysis.

### EEG analysis reaching task

We performed the same preprocessing and time-frequency transformation steps as described in the fist clenching task. The data was segmented into epochs from −1.5 to 3 s relative to the cue color change, i.e., the *go*-signal to imagine the reach to the middle target. After the time-frequency transformation, we cropped the TFR data from −1 to 2.5 s and set the baseline window to −0.75 to −0.25 s in relation to the start of the imagined reach.

#### Correlation analysis imagined fist clenching task

To relate the neural data of the imagined fist clenching task to performance in the reaching task, we combined MPE change adaptation, MPE change baseline/washout and FFC final adaptation to one behavioral value. Our aim was to obtain one behavioral measure that reliably captured overall motor adaptation performance. For this, we normalized each dependent variable by dividing it by its standard deviation across participants. Next, we added MPE change adaptation, MPE change baseline/washout and the inverse of FFC final adaptation. The resulting behavioral variable was termed change of error. A more negative value denoted a better overall adaptation performance.

We performed a cluster-based permutation test to investigate any possible association between the behavioral change of error and the neural data of the imagined fist clenching task. For this, we first correlated change of error with the calculated TFR value for each time-frequency-channel bin. We set the initial threshold for each correlation to *p* = 0.01. Directly neighboring significant bins in time, frequency, or channel formed a cluster. We summed the t-values of those bins belonging to the same cluster and recorded the biggest absolute value of the sum. To obtain an empirical distribution we permuted change of error randomly across participants and repeated the described steps 1000 times. Finally, we compared the originally observed biggest cluster mass to the empirical distribution. Here, we used the standard p-value cutoff 0.05 (see Fig. 4A & B; note that due to thresholding, correlation values are partially underestimated in 4B).

In addition, to account for different time courses in the neural signature of participants, we calculated the correlation of change of error and the largest positive (ERS) and largest negative (ERD) power change value in channel C3. We picked C3 because it is over the primary motor cortex, contralateral to the imagined movement side, and therefore, is usually investigated in motor imagery research as the sensor most reliably picking up activity from the motor cortex (e.g., ^5^). Since we observed the greatest power modulation averaged over participants (see Fig. 3C) in alpha band, we took the maximal and minimal power values between 8 and 13 Hz. Also based on the average strength of ERD and ERS, we selected the peak ERD value in a time window between 0 and 2 s and the peak ERS value in a time window between 2 and 4 s. Lastly, we also calculated the correlation of change of error and the difference between peak ERS and ERD. We applied the Bonferroni-Holm correction to adjust the p-values of the 3 correlation tests.

To illustrate the correlation of the strength of ERS with adaptation performance in source space, we first averaged the power change of each participant over the ERS time window (2.5 - 3s) and the alpha frequency band (8 - 13Hz) for all sources. Next, we correlated the average power change of each source with the change of error in the adaptation task across participants (see Fig. 4D). We performed the same calculation steps for the subsequent time window (3 - 3.5s; see Fig. 4E). To observe the correlation strength over different time and frequency ranges in source space, we also plotted the average correlation of 1s segments in the examined frequency ranges (see supplementary Fig. 3B, C, D).

#### Correlation analysis reaching task

We performed an analogous analysis to relate the neural data of the reaching task to the change of error in the same task.

#### Correlation analysis questionnaire

Lastly, we investigated if there was any association between the self-reported ease/difficulty of imagining the feeling of the movement in the reaching task and the degree of motor adaptation. For this, we correlated the subjective imagery rating with the change of error across participants. In addition, we repeated the described cluster-based permutation tests above but replaced the change of error with the subjective imagery rating. Again, we set the initial threshold to 0.01. However, we only permuted the data 100 times to save resources as it was evident that significance would not be reached in these analyses. Finally, we correlated the subjective imagery rating with the peak ERS, peak ERD, and the difference between peak ERS and ERD in both tasks.

### 4.5 Preregistration

We preregistered our study design of the reaching task, which includes sample sizes, general hypotheses, and the main kinematic analysis plan, on OSF (https://osf.io/swgd9). Furthermore, exemplary reaching task videos are available there.

### 4.6 Data availability

Kinematic data and EEG data of the imagined fist clenching task will be publicly available on OSF upon publication.

### 4.7 Code availability

The code used for data processing, analysis, and visualization in this study will be publicly available on GitHub upon publication.

### 4.9 Author contributions

MG, TH, VN and BS conceived the study. MG and P-CS programmed the tasks. MG acquired and analyzed the data. MJI and VN advised on EEG processing. MJI reviewed the code. MG wrote the original draft of the manuscript. All authors reviewed and contributed to the manuscript.

## Acknowledgements

The authors thank Elisabeth Kaminski and Philipp Kaniuth for valuable discussion, Joshua Grant for proof-reading, and Lisa Franke and Saskia Leupold for help with data collection.

