## Supplementary Information for "Motor Imagery Enhances Performance Beyond the Imagined Action"

### 5 Supplementary Information

#### 5.1 Supplementary Results 1

**Force field compensation comparison across and within groups.**

Comparing the average FFC value at the end of the adaptation phase against 0 revealed significant within-group effects for the active ($t(19) = 27.04$, $p_{active} = 6.3\text{e-}16$) and MI group ($t(19) = 9{,}16$, $p_{MI} = 6.3\text{e-}08$) but not for the control group ($t(19) = 2.22$, $p_{control} = 0.039$; see Fig. 2F). Similarly, t-tests between groups revealed differences between all groups ($t(18) = 17.02$, $p_{active/control} = 1.4\text{e-}18$; $t(18) = 7.64$, $p_{active/MI} = 1.4\text{e-}08$; $t(18) = 6.16$, $p_{MI/control} = 6.9\text{e-}07$). The active group showed the most predictive compensation, followed by the MI group.

#### 5.2 Supplementary Results 2

**Power changes during the imagined fist clenching and reaching task.**

On a descriptive level the ERD was strongest in the alpha band (8-13 Hz; see Fig. 3B) over the contralateral left hemisphere (see Fig. 3C). We also observed an event-related synchronization (ERS) at the end of the motor imagery phase which, however, showed a weaker lateralization. Source reconstruction of the ERD in the alpha frequency band revealed that the strongest desynchronization, according to the Desikan-Killiany atlas, was observed in the left post- and precentral region, i.e. the primary sensory and motor cortex, respectively (see Fig. 3D, left). The ERS was also present in the contralateral hemisphere and more pronounced in the posterior parts of the cortex. The synchronization was strongest in the left superior-parietal region (see Fig. 3D, right). In beta frequency bands, topoplots of the ERD and ERS and the respective source reconstructions showed a similar pattern (see supplementary Fig. 1A & B).

The activity pattern over sensorimotor areas during the reaching task showed lateralization of an ERD in the alpha band during the overt reach with a stronger desynchronization on the ipsilateral, right side. The ERD was similar in strength on both hemispheres in the beta band. In addition, there was an ERS during the imagined reach that was stronger on the contralateral, left side (see supplementary Fig. 2A & B).

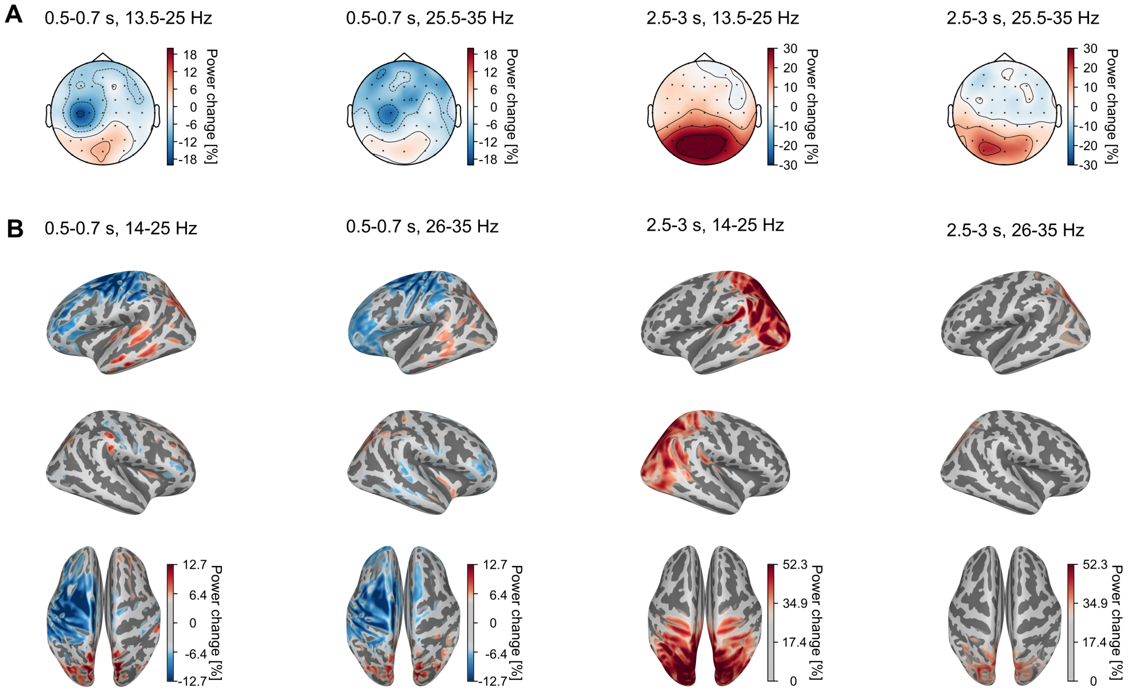


Supplementary Figure 1: **EEG data imagined fist clenching task (beta frequencies.)** Wherever possible, the color bars were maintained identical to those in Fig. 3 to facilitate comparison. A) Topoplots of averaged beta and high beta activity in chosen time windows averaged across participants. B) Source reconstructed inflated brains of averaged beta activity in chosen time windows averaged across participants.

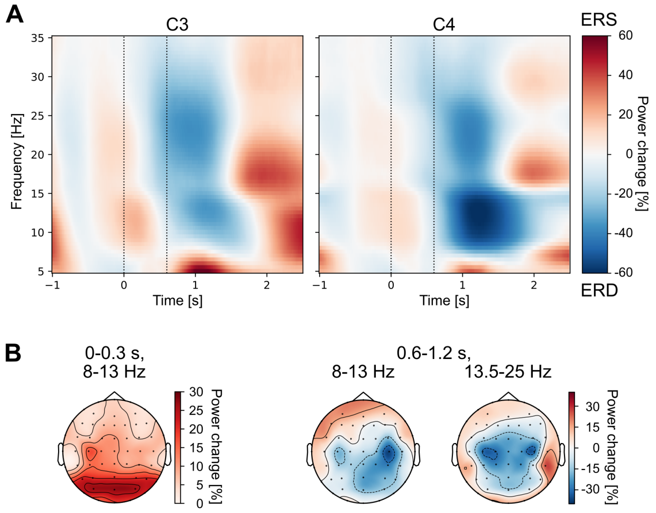


Supplementary Figure 2: **EEG data reaching task.**

A) Time-frequency representation of grand averaged (n = 18) data in C3 and C4. Color represents power change in percent relative to the baseline window from -0.75 to -0.25s. The *go*-signals for the imagined and overt reach were at 0 and 0.6 s, respectively. B) Topoplots of averaged alpha and beta activity in chosen time windows averaged across participants.

#### 5.3 Supplementary Results 3

**Additional visualizations of correlations between change of error in the reaching task and oscillatory power changes in the imagined fist clenching task.**

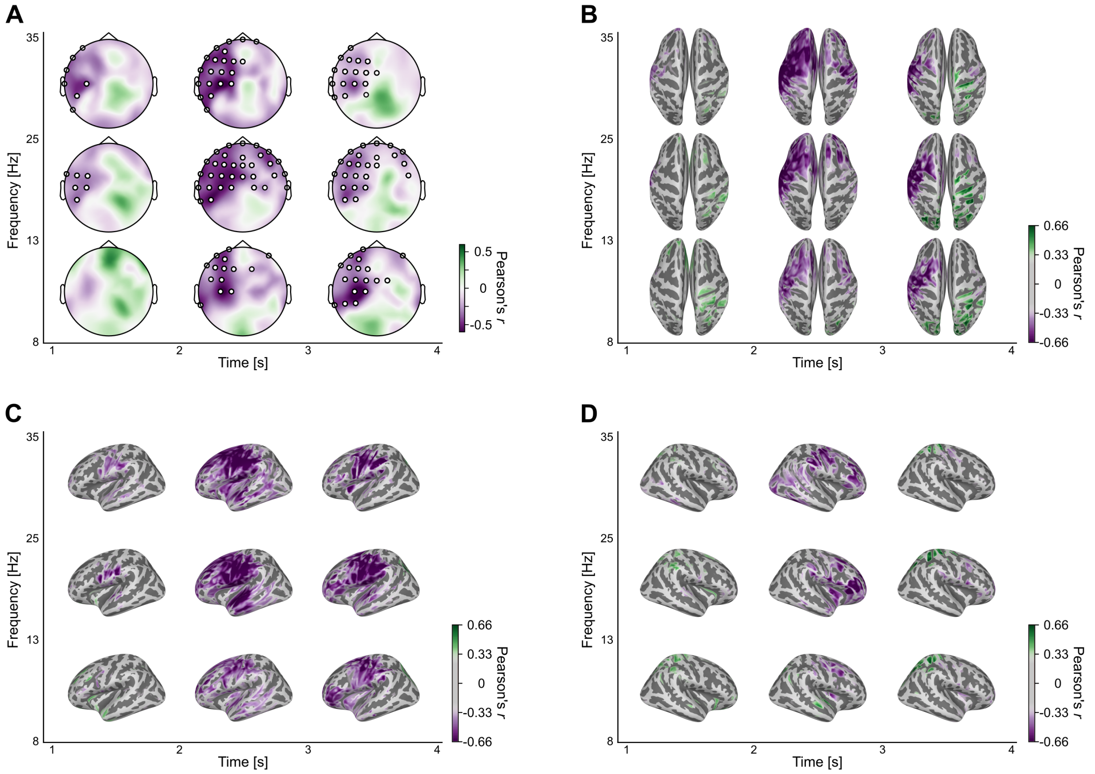


Supplementary Figure 3: **Pearson's correlations of change of error in the reaching task with power change in the imagined fist clenching task across participants.** A) Topoplots of different time windows and frequency ranges displaying the distribution of correlation values. Correlation values were averaged in the specified time and frequency ranges. Channels of the significant cluster are marked in white. B), C) & D) Inflated brains display correlation values for different time windows and frequency ranges in source space. Correlation values are averaged in the specified time and frequency ranges. The colorbars were maintained identical to those in Fig. 4D to facilitate comparison. B) Dorsal view. C) Lateral view left hemisphere. D) Lateral view right hemisphere.

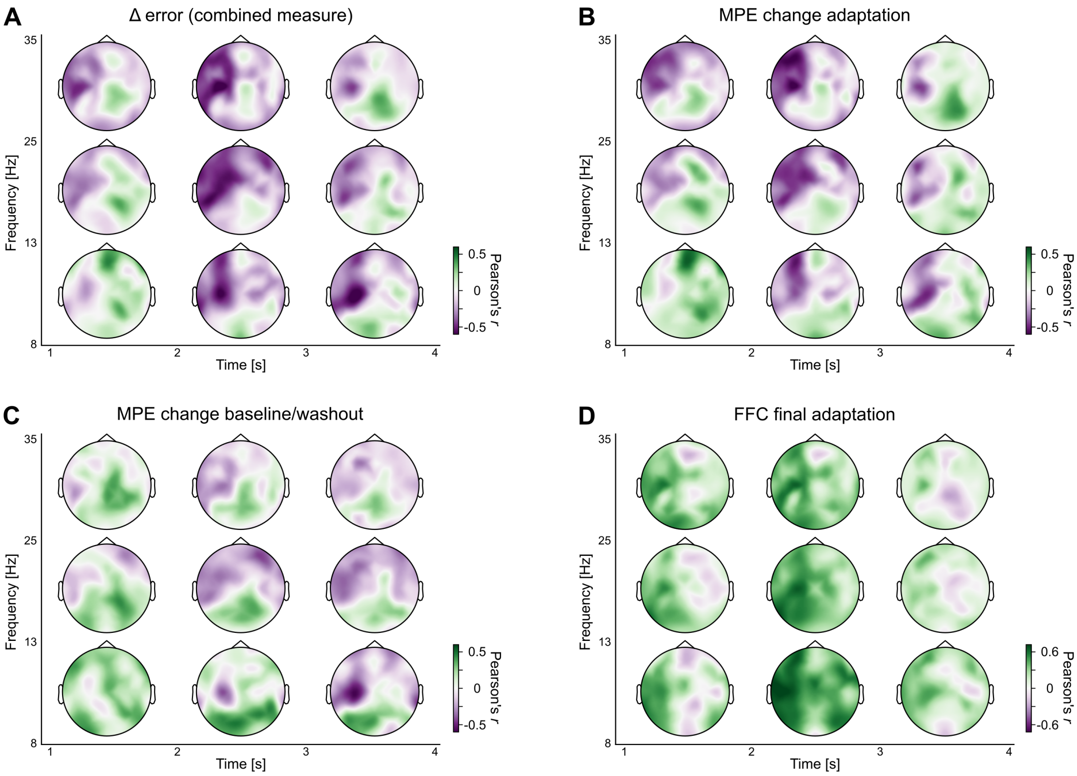


Supplementary Figure 4: **Pearson's correlations of behavioral measures in the reaching task with power change in the imagined fist clenching task across participants.** Topoplots of different time windows and frequency ranges displaying the distribution of correlation values. Correlation values were averaged in the specified time and frequency ranges. We have shown before that behavioral measures are strongly correlated in our specific setup of this task[26]. A) Combined measure (change of error) for comparison. B) Change of MPE from beginning to end of adaptation. C) Change of MPE from baseline to washout. D) Average FFC in the last 4 adaptation blocks. Note that here the inverse of the correlation pattern is expected, because the better the performance improvement, the bigger the FFC final adaptation measure.

Table 1: **Relationship between behavior and EEG measures.** Pearson's correlation of change of error in the reaching task with peak power changes in the imagined fist clenching task across participants in C3 for different scenarios. ERS - maximal positive power change value in the alpha band (if not indicated differently) between 2 and 4s. ERD - maximal negative power change value in the alpha band (if not indicated differently) between 0 and 2s. Rebound - difference between peak ERS and peak ERD. All p-values are uncorrected. Significant correlations are in bold.

| Change of error correlated with | ERS | ERD | Rebound |
|---|---|---|---|
| Exclusion of participant #51 (most influential for ERS correlation) | $r(13)$ = **-0.615** $p$ = 0.015 | $r(13)$ = 0.156 $p$ = 0.579 | $r(13)$ = **-0.682** $p$ = 0.005 |
| Exclusion of participant #29 (most influential for rebound correlation) | $r(13)$ = **-0.676** $p$ = 0.006 | $r(13)$ = -0.282 $p$ = 0.309 | $r(13)$ = **-0.674** $p$ = 0.006 |
| Baseline window -1 : 0 (instead of -0.75 : -0.25) | $r(14)$ = **-0.764** $p$ < 0.001 | $r(14)$ = -0.122 $p$ = 0.653 | $r(14)$ = **-0.726** $p$ = 0.001 |
| Average reference (instead of current source density) | $r(14)$ = **-0.52** $p$ = 0.039 | $r(14)$ = 0.136 $p$ = 0.616 | $r(14)$ = **-0.574** $p$ = 0.02 |
| Morlet wavelet transformation (instead of multitaper) | $r(14)$ = **-0.679** $p$ = 0.004 | $r(14)$ = 0.078 $p$ = 0.774 | $r(14)$ = **-0.687** $p$ = 0.003 |
| Beta band 13.5 - 25 Hz (instead of alpha 8 - 13 Hz) | $r(14)$ = **-0.634** $p$ = 0.008 | $r(14)$ = 0.046 $p$ = 0.866 | $r(14)$ = **-0.662** $p$ = 0.005 |
| High beta band 25.5 - 35 Hz (instead of alpha 8 - 13 Hz) | $r(14)$ = **-0.551** $p$ = 0.027 | $r(14)$ = -0.391 $p$ = 0.134 | $r(14)$ = -0.354 $p$ = 0.179 |

#### 5.4 Supplementary Results 4

**Reaching task performance is not related to neural data during the same task.**

We did not find a relationship between change of error and average power change in the reaching task in a cluster-based permutation test ($p$ = 0.22). Similarly, we also did not find a significant correlation between adaptation performance and peak ERD ($r(16)$ = 0.177, $p$ =0.482), ERS ($r(16)$ = 0.156, $p$ = 0.536) or peak-to-peak difference ($r(16)$ = -0.019, $p$ = 0.94) in the 600 ms motor imagery time window in alpha band across participants. This lack of significant findings remained the same when we related neural activity in the last 100 trials to adaptation performance, and also when we segmented the EEG data relative to actual movement onset and not to the visual cues indicating the start of the (imaginary) movement. Taken together, we could not find a neural marker during motor imagery in the reaching task predicting the overall motor adaptation performance of participants.

#### 5.5 Supplementary Results 5

**Subjective rating of motor imagery is not related to performance or neural data.**

To investigate if there was a relationship between perceived proficiency in motor imagery and motor performance or neural data, we asked participants in the MI group about their subjective experience at the end of the experiment. We adjusted questions from a widely used motor imagery questionnaire (MIQ-RS,[51]) to fit our task. In particular, we asked participants to indicate how successfully they were able to feel the imagined prior movement on a Likert Scale from 1 ("very hard to feel") to 7 ("very easy to feel") on average throughout the reaching task. If self-assessed motor imagery would correctly characterize the actual ability to perform motor imagery, we would expect to see a relationship between this subjective imagery rating and the degree of adaptation in the reaching task. However, there was no correlation between the imagery rating and change of error in the reaching task across participants ($r(18)$ = -0.030, $p$ = 0.91).

We also investigated the relationship between the subjective imagery rating of the reaching task and the neural response in the imagined fist clenching and reaching task, respectively. We adopted the same approach as before when we related neural data and change of error. We did not find any significant clusters when performing a cluster-based permutation test over the complete window duration and considered frequency ranges in the imagined fist clenching ($p$ = 0.25) or reaching task (0.96). We also did not find any relationship between the imagery rating and peak ERS, imagery rating and peak ERD, or imagery rating and peak-to-peak difference of ERS and ERD in the alpha band in the imagined fist clenching (ERS: $r(14)$ = 0.33, $p$ = 0.21; ERD: $r(14)$ = 0.35, $p$ = 0.18; ERS - ERD: $r(14)$ = 0.16, $p$ = 0.55) or in the reaching task (ERS: $r(16)$ = 0.09, $p$ = 0.722; ERD: $r(16)$ = 0.279, $p$ = 0.262; ERS - ERD: $r(16)$ = -0.214, $p$ = 0.394). Taken together, we did not find any evidence that the subjective rating of motor imagery difficulty/ease is related to performance or a neural manifestation.
